# Ecological assessment of the River Nile around Gizert El-Warrak by Phytoplankton and macroinvertebrates assemblages

**DOI:** 10.1101/297432

**Authors:** Kadria M.A. Mahmoud, Sara S.M. Sayed, Mohamed R. Habib

## Abstract

Biological assessment is considered a useful tool for assessing the ecological status of the aquatic ecosystems. Therefore, the goal of the present study was to use phytoplankton and macroinvertebrates as biological tools for ecological assessment of the River Nile around Gizert El-Warrak. A total of 33 phytoplankton species identified in the studied stations; these include 16 species of Chlorophyta, 5 species of Cyanophyta and 12 species of Bacillariophyta. All investigated stations characterized by high organic pollution according to Palmer′s index. Trophic state index showed a hyper-eutrophic status in stations S1, S2, S4, S6 and S8 and an eutrophic status in stations S3, S5 and S7.Gastropoda and Oligochaeta were the most dominant of macroinvertebrates taxa recorded 50.8 and 24.6%, respectively. Diversity Index (H′) ranged (1.14 – 2) which indicated that the structure of macroinvertebrates habitat was poor. Also, Evenness Index (J) ranged (0.016 – 0.043) which indicated that individuals were not distributed equally. The values of biotic index depending on macroinvertebrates categories showed that the River Nile’s water quality is fairly poor with significant organic pollution.

**Summary statement:** Bioassessment based on non-taxonomic measurements of algae and biotic indices of macroinvertebrates may be considered as vital methods that reflect disturbances in aquatic systems for both short-term and long-term.

## Introduction

Although the chemical analysis of water gives a good sign of the quality of the aquatic system, it does not necessarily reflect the ecological status of the system (Karr et al., 2000). Generally, the chemical assessment of water quality is based on determining the most important water parameters; nitrate, nitrite, soluble reactive phosphorus, ammonia, oxygen and biochemical oxygen demand. Whereas phosphorus and nitrogen are the relevant parameters for assessing the nutrient loading, ammonia and o xygen saturation are the pertinent criteria with respect to saprobity levels which demonstrated that these environments are undergoing a process of degradation (Rangel et al., 2012). Nevertheless, the most acceptable ecosystem assessment requires evaluating physical and chemical factors as well as the composition and structure of biotic assemblages (Lobo and Callegaro, 2000).

In the last few decades, several developed and developing countries have been interested in rapid evaluation methods for the bioassessment of water quality (Al-Shami et al., 2011). Most of these methods depend on phytoplankton and macroinvertebrates for the development of adequate tools to measure the ecological status of freshwater systems. In addition, these communities give insights about both the environmental effects of water chemistry and the physical characteristics of rivers (Stevenson and Pan, 1999). Furthermore, bioindication is considered easy and cost-effective tools for short- and long-term monitoring of environmental and ecosystem integrity (Neumann et al., 2003).

Phytoplankton is considered as a suitable bioindicator, they have a worldwide distribution, high reproduction rate and each species have high sensitivity towards different levels of organically polluted waters (Moura et al., 2007). Also, they are used as indicators for saprobic conditions such as salinity, acidification and eutrophication in lakes and rivers (Smith et al., 2014). Moreover, pH, ionic strength, substrate, current velocity, light (degree of shading), grazing and temperature affect the distribution patterns of phytoplankton in lotic systems (Santos et al., 2016).). Also, Chlorophyll-*a* concentrations have been a subject of interest for many researchers concerned with the water quality for determining phytoplankton distribution as an indicator of the water body’s health, composition, and ecological status (Kasprazak et al., 2008). At water surface, high chlorophyll-*a* levels indicated to high algal growth or blooming and this is usually associated with excessive nutrients such as phosphorus and nitrogen. Upon dying, algae are depleted dissolved oxygen levels and lead to fish death (Horrigan et al., 2002).

Another approach in water quality assessment, it is macroinvertebrates assemblages that have been traditionally used in the biomonitoring of stream and river ecosystems for various environmental stress types, such as organic pollution (Li et al., 2010) and river health (Sharifinia et al., 2016). Macroinvertebrates-based biotic index has been developed for rapid bio-assessment of rivers (Elias et al., 2014; Kaaya et al., 2015), since they are adapted to specific environmental conditions. If these conditions change, some organisms can disappear and be replaced by others. Therefore, variations in the composition and structure of their assemblages in running waters can indicate possible pollution (Alba-Tercedor, 1996). Gizert El-Warrak is an Egyptian island in the River Nile, highly populated and many human activities are conducted, a huge amount of sewage and drainage waters are discharged in the River Nile affecting many aquatic organisms (Sleem and Hassan, 2010). However, there was no more study done about the River Nile’s water quality around Gizert El-Warrak. So that, using bioindication depending on phytoplankton and macroinvertebrates assemblages may give us a scope to assess the ecological status of the River Nile water quality around Gizert El-Warrak in the current work that aims at (1) estimate chlorophyll-*a* content, ammonia, nitrite, nitrate and phosphate for the assessment of the present water quality in the River Nile, (2) apply different indices based on phytoplankton and macroinvertebrates for determining water pollution levels and (3) study the compatibility of biological and chemical results to give an integrative picture about the water quality of the River Nile.

## Material and Methods

### Study area

Gizert El-Warrak is an Egyptian island in the River Nile with an area of about 1400 acres. The island located in El-Warrak city in Giza governorate, Egypt. It has a distinguishable location, bordered by Qalyubiya governorate from the North and Cairo governorate from the East. The study was conducted during December 2016 to February 2017 in the River Nile around Gizert El-Warrak from two sides of the island using a boat. Four stations were selected from each side, at a distance about 100 meters between stations. Stations, S1-S4 lies at 30º7′6.99″N which toward Warrak EL-Hadder city and the main human activities were farming practices, while stations S5-S8 lies at 31º13′32″E which toward Shubra city and fishing was the common activities **(Fig. 1).**

**Figure 1.**
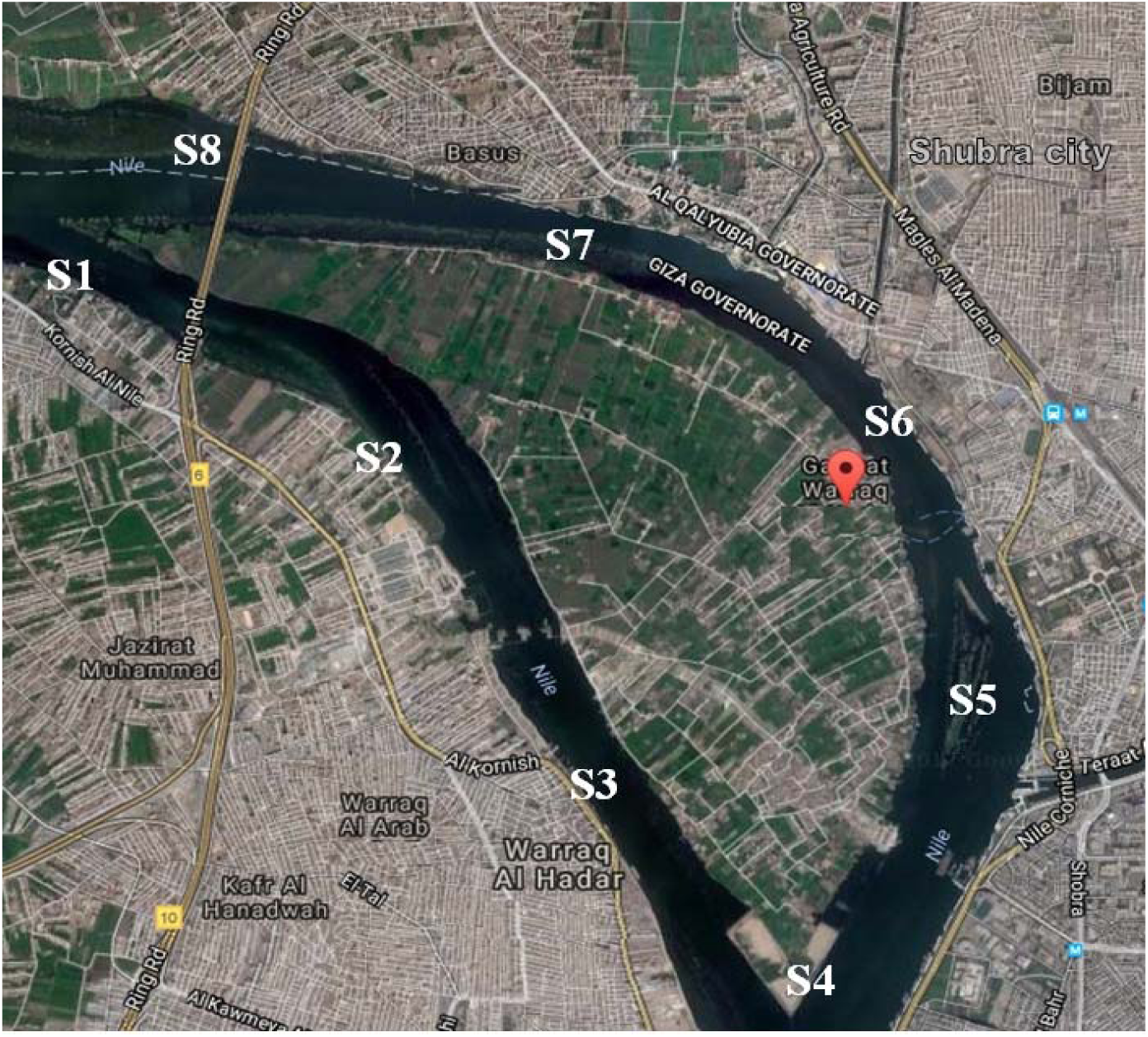
A map illustrated the location of Gizert El-Warrak and the investigated stations.

### Field study

Total dissolved solids (TDS), water temperature (ºC) and electrical conductivity (EC) were measured monthly using a portable conductivity meter (HI 9635) and a portable pH meter (HI 9024) in three times per each station *in situ* at a mid-day, at 20 cm under the water surfaces. About three liters of water samples from each station were filtered *in situ* through a plankton net with mesh diameter 20μm and specimen were preserved by adding few drops of 4% formaldehyde. All samples were transferred in the labeled plastic containers to the laboratory for identification. Macroinvertebrates samples were collected in five times, using D-shaped aquatic net (20 × 40 cm) equipped with about 1m. long handle. Additionally, all macrophytes were removed from the stream and visually searched for macroinvertebrates, then all samples were transferred in the labeled plastic containers to the laboratory for identification.

### Laboratory study

Water samples were collected from each station in three liters’ plastic containers by the simple dip method and transferred to the laboratory. Nitrite (NO_2_D□), nitrate (NO_3_D□), ammonia (NH_3_^+^), phosphate and quantification of chlorophyll-*a* concentration were determined in collected water samples by standard methods according to APHA (2005).

The taxonomic composition of the phytoplankton was determined by a microscope Olympus 200x according to common taxonomic keys stated by Komárek and Komarkova (1992).

The trophic status of each station was calculated according to Carlson (1977), depending on chlorophyll-a concentration using the following equation:

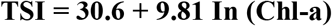

Where, TSI = Trophic State Index, ln = natural logarithm and (Chl-a) = concentration of chlorophyll-a (μg/l). TSI value indicated as follows: >30 = Ultra oligotrophic, 30-50 = Oligotrophic, 51-60 = Mesotrophic, 61-70 = Eutrophic and <70 = Hyper-eutrophic.

On the other hand, organic pollution/ site was evaluated according to the Palmer′s pollution index (Palmer, 1968) depending on the major list of the algal genera that are tolerant to organic pollution. Palmer′s pollution index score was indicated as follows: <15 = very light organic pollution, 16-20 = moderate organic pollution and >20 = high organic pollution.

Macroinvertebrates samples were washed with dechlorinated tap water through a sieve (300 mm pore diameter), sorted out and identified according to published keys (Hynes, 1984; Elliott et al., 1988; Pescador et al., 2004).

Diversity Index (H′) was used according to Shannon-Wiener (1949) formula:

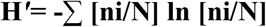

Where **ni** is the number of individuals in each species, **N** equals the total number of individuals in the sample, and ∑ equals the total number of species in the sample. Results are generally between >4 = high status, 4-3= good status, 2-1= poor status and <1 =bad status.

Evenness Index (J) was used according to Pielou (1966) formula: **J = H′/ In S**

H = Shannon – Wiener diversity index, ln = natural logarithm and S = total number of species in the sample. The values are between 0 – 1.When the value is getting closer to 1, it means that the individuals are distributed equally.

Biotic Index (BI) is based on categorizing macroinvertebrates into categories depending on their response to organic pollution according to Hilsenhoff (1977) formula: **BI =** ∑ **[(ni *ai)/N]**

Where **ni** is the number of specimens in each taxonomic group, **ai** is the pollution tolerance score for that taxonomic group, and **N** is the total number of organisms in sample.

### Statistical analysis

Correlation between physicochemical variables and macroinvertebrates were described by correspondence analysis (CA). The abundance of macroinvertebrates orders was performed by Box plot using XLSTAT 2016, Statistical software for Microsoft Excel, Par-is, France.

## Results and Discussion

The average values of water temperature, pH, electrical conductivity (EC), total dissolved solids (TDS), ammonia (NH_3_^+^), nitrite (NO_2_D□), nitrate (NO_3_D□) and phosphate (PO_4_) are shown in **Table (1)**. Generally, pH range of the present water samples was alkaline (8.5-9). These results were in parallel to Svobodova et al. (1993) who declared that high water alkalinity was attributed to uptake a considerable amount of CO_2_ during the day by algae and aquatic plants for photosynthetic activity in eutrophic waters. Phosphate concentrations at stations S1, S2, S3, S4, S5 and S6 were higher than stations S7 and S8. The highest concentrations of ammonia (303 mg/l) and nitrate (271 mg/l) were recorded at stations S2. This may be due to runoff of agriculture wastewater during farming practices. Yadav and Rajesh (2011) declared that phosphate levels increased during winter due to agricultural runoff containing phosphate fertilizers and (detergents) municipal wastewater. Nevertheless, high concentrations of phosphate are rarely found in water where it is actively consumed by aquatic plants. In addition, TSI results indicated that stations S1, S2, S4, S6 and S8 were hyper-eutrophic and water quality was poor, while stations S3, S5 and S7 were eutrophic and water quality was fair as shown in **Table (2)**. These results may attributed to high chlorophyll-*a* content which is often used as an estimate of algal biomass, these results were in coincidence with the pervious studied by Wetzel (2001).

**Table 1.** The mean of physico-chemical characteristics of water collected from the eight investigated stations.

| Stations | T<br>(°C ) | EC<br>(µs/cm) | pH | TDS<br>(mg/l) | Ammonia<br>(NH<sub>3</sub>)<br>(µg/l) | Nitrite<br>(NO<sub>2</sub>)<br>(µg/l) | Nitrate<br>(NO<sub>3</sub>)<br>(µg/l) | Phosphate<br>(PO<sub>4</sub>)<br>(µg/l) |
| --- | --- | --- | --- | --- | --- | --- | --- | --- |
| S1 | 18.9±0.51 | 460±0.04 | 8.5±0.17 | 220±0.02 | 49±0.04 | 38±0.01 | 175±0.05 | 114±0.05 |
| S2 | 19.3±1.34 | 430±0.03 | 8.7±0.16 | 280±0.10 | 303±0.36 | 25±0.02 | 271±0.07 | 167±0.08 |
| S3 | 18.7±1.70 | 440±0.04 | 8.7±0.20 | 210±0.03 | 49±0.04 | 23±0.02 | 153±0.06 | 104±0.06 |
| S4 | 18.0±1.75 | 360±0.09 | 8.8±0.15 | 210±0.03 | 60±0.06 | 51±0.03 | 201±0.01 | 151±0.08 |
| S5 | 18.2±0.76 | 450±0.05 | 9.0±0.05 | 222±0.02 | 33±0.06 | 44±0.01 | 181±0.09 | 124±0.05 |
| S6 | 18.2±1.60 | 420±0.07 | 8.9±0.12 | 220±0.02 | ND | 40±0.02 | 206±0.10 | 248±0.25 |
| S7 | 18.0±1.00 | 450±0.02 | 8.9±0.04 | 230±0.01 | ND | 30±0.01 | 172±0.10 | 83±0.02 |
| S8 | 18.3±0.43 | 440±0.03 | 9.0±0.06 | 220±0.01 | ND | 38±0.02 | 253±0.10 | 48±0.04 |

Temperature (T), Electrical conductivity (EC) and Total dissolved solids (TDS), Not Detected (ND).

**Table 2.** List of the identified phytoplankton taxa and scoring of Palmer’s pollution index of the River Nile at Gizert El-Warrak stations.

| Carlson's Trophic state index (TSI) | Criteria | sites | Mean of Chl-*a*<br>(µg/l) | TSI calculated/<br>site | Trophic state/site | Water quality depend on TSI/ site |
| --- | --- | --- | --- | --- | --- | --- |
| Ultraoligotrophic | <30 | S1 | 170.5 | 81.01 | Hyper-eutrophic | Poor |
| Oligotrophic | 30-50 | S2 | 150.2 | 79.76 | Hyper-eutrophic | Poor |
| Mesotrophic | 51-60 | S3 | 30.26 | 64.05 | Eutrophic | Fair |
| Eutrophic | 61-70 | S4 | 95.13 | 75.28 | Hyper-eutrophic | Poor |
| Hyper-eutrophic | >70 | S5 | 34.83 | 65.43 | Eutrophic | Fair |
| Water quality depend on TSI values |  | S6 | 62.50 | 71.16 | Hyper-eutrophic | Poor |
| Good | 0-59 | S7 | 40.13 | 66.81 | Eutrophic | Fair |
| Fair | 60-69 | S8 | 66.56 | 71.78 | Hyper-eutrophic | Poor |
| Poor | 70-100 |  |  |  |  |  |

In the current work, a total of 33 identified phytoplankton species were 16 species of Chlorophyta (green algae); 5 species of Cyanophyta (blue green algae) and 12 species of Bacillariophyta (diatoms). Diatoms were more dominant than green algae and blue green algae which showed an appreciable presentation **(Table 3)**. Bilous et al. (2012) reflected that the eutrophic state of the river ecosystem mostly due to the presence of green algae and diatoms. These results were agreed with Ganai and Parveen (2014) who stated that Bacillariophyta was the most dominant group of phytoplankton community in Wular Lake at Lankrishipora, Kashmir, and attributed that to their ability to grow under unsuitable conditions such as weak light, low temperature and nutrients (Ganai et al., 2010). In present work, all the investigated stations characterized by high organic pollution according to palmer’s index **(Table 3)**, that may explain the low number of the identified phytoplankton taxa as many of species might be disappeared due to the heavy pollution. The green algae (*Actinastrum* sp., *Ankistrodesmus acicularis, Scenedesmus obliquus* and *Scenedesmus quadricauda*) and diatoms (*Diatoma elongatum, Syndra ulna, Nitzschia acicularis, Nitzschia filiformis* and *Nitzschia liniaris*) were the most dominant in the present examined stations. These findings are in agree with Szabo et al. (2005) who demonstrated that *Achnantidium minutissimum* and many *Nitzschia* species are tolerant to a variety of pollution and are dominated in the environments that have stress. Furthermore, Kumar and Sharma (2014) explained that some species such as *Aulacoseria granulate, Cocconeis placentula, Cymbella spp., Fragilaria capucina, Gomphonies olivaceum, Diatoma elongatum, Navicula radiosa* and *Syndra ulna* were indicative to trophic status of aquatic ecosystems between oligotrophic to eutrophic

**Table 3.** Classification of the eight investigated stations according to the Carlson’s Trophic State Index (TSI) depending on average chlorophyll-*a* concentration.

|  | Algae Taxa | S1 | S2 | S3 | S4 | S5 | S6 | S7 | S8 |
| --- | --- | --- | --- | --- | --- | --- | --- | --- | --- |
| Chlorophyta | *Actinastrum sp.* | + |  | + |  | + | + | + | + |
|  | ***Ankistrodesmus acicularis*** | + (2) | +(2) | +(2) | +(2) | +(2) | +(2) | ++(2) | ++(2) |
|  | *Chodatella cillata* | + | + |  | + |  |  |  |  |
|  | *Coelastrum microporum* | + | + | + |  | + |  |  | + |
|  | *Cosmarium bioculatum* | ++ | + | + |  |  |  |  |  |
|  | *Mougeotia scalaris* | ± | + |  | + |  |  | + | +(1) |
|  | *Nephrocytium sp.* | + |  | + |  |  |  |  |  |
|  | *Oocystis parva* | + |  |  |  |  | + | + |  |
|  | *Pediastrum clathratum* | + |  | + | + |  |  |  |  |
|  | *Pediastrum gracilimum* |  | + |  |  | + | + |  | + |
|  | *Pediastrum tetras* |  | + |  |  |  | + | + |  |
|  | ***Scenedesmus obliquus*** | +(4) |  | +(4) | +(4) |  |  |  |  |
|  | ***Scenedesmus quadricauda*** | ++(4) | +(4) | +(4) | ++(4) | +(4) | +(4) | +(4) | ++(4) |
|  | *Staurastrum paradoxum* | + | + |  |  | +(4) | +(4) |  | +(4) |
|  | *Tetraedron minimum* | + | + | + | + | + | + | + | ++ |
|  | *Ulothrix subtilissima* | + |  | + |  | + | + | + |  |
| Cyanophyta | ***Gomphasperia lacustris*** | +(11) | +(1) |  | +(4) | +(1) | +(1) | +(1) | +(1) |
|  | *Merismopedia elegans* | + | + | + | + | + | + | + | + |
|  | ***Phormidium sp.*** | +(1) | +(1) |  |  |  |  |  |  |
|  | ***Oscillatoria limnetica*** |  | +(4) | +(4) | + | +(4) | +(4) | +(4) | +(4) |
|  | *Microcystis flos-aquae* | ± | + | + |  |  | + | + | + |
| Bacillariophyta | *Amphora coffeaeformis* |  | + |  |  | + |  |  | + |
|  | *Cocconies placentula* | + | + |  |  |  |  |  | + |
|  | ***Cyclotella comta*** | ++(1) | ++(1) | ++(1) | ++(1) | +++ (1) | +++ (1) | +++ (1) | +++ (1) |
|  | *Diatoma elongatum* | +++ | +++ | ++ | + | +++ | +++ | +++ | +++ |
|  | *Fragilaria capucina* | +++ | +++ | + | ++ |  |  |  |  |
|  | ***Melosira granulata*** | ++(1) | ++(1) | +(1) | +(1) | ++(1) | ++(1) | ++(1) | ++(1) |
|  | *Navicula cuspidata* | +(3) | +(3) |  |  | +(3) |  | +(3) |  |
|  | ***Nitzschia acicularis*** | ++(3) | +++ (3) | +(3) | ++(3) | +(3) | +(3) |  | +(3) |
|  | ***Nitzschia filiformis*** | ++(3) | ++(3) |  |  | +(3) | +(3) | +(3) | +(3) |
|  | ***Nitzschia liniaris*** | ++(3) | +(3) |  | +(3) | ++(3) | +(3) | +(3) | +(3) |
|  | *Stephanodiscus dubius* | ++ | + |  | + | + |  | + | + |
|  | ***Synedra ulna*** | +(2) | ++(2) | +(2) | +(2) | ++(2) | ++(2) | ++(2) | ++(2) |
|  | **Palmer's pollution index** | **38** | **28** | **21** | **24** | **29** | **31** | **28** | **24** |

++++: Dominant; +++: Plenty; ++: Many; +: Appreciable; ±: Rare. Species in bold font represent major list of algal genera that tolerant to organic pollution and their scoring. Palmer's pollution index: <15 very light organic pollution, 16-20 moderate organic pollution and >20 high organic pollution.

Data of the collected macroinvertebrate taxa from sampling stations are presented in **Table (4)**. Generally, the highest number of taxa was recorded at S1 and S4 with 789 and 776, respectively. Gastropoda and Oligochaeta were the highest taxa in relative abundance **(Fig. 2)**. This is in parallel to the pervious study of El-Khayat et al. (2011) who indicted that freshwater snails are generally tolerant to organic pollution. In addition, Andem et al. (2015) stated that the highest oligocheata abundance is an indication to poor water quality of Ediba River in Cross River State, Nigeria. Hence, a high density of them is a good indication of organic pollution because they are able to tolerate unfavorable conditions such as low dissolved oxygen and high pollutant concentrations.

**Table 4.** Taxonomic composition of macroinvertebrates taxa of the River Nile at Gizert El-Warrak stations.

|  | Macroinvertebrates orders | Code | Taxon | S1 | S2 | S3 | S4 | S5 | S6 | S7 | S8 | Total | % |
| --- | --- | --- | --- | --- | --- | --- | --- | --- | --- | --- | --- | --- | --- |
| Insecta macroinvertebrates | Ephemeroptera | Eph | Mayflies | 21 | 19 | 9 | 47 | 13 | 8 | 26 | 35 | 178 | 3.86 |
|  | Plecoptera | Pleo | Stoneflies | 12 | 12 | 10 | 12 | 12 |  | 3 | 7 | 68 | 1.47 |
|  | Diptera (Chironomidae) | Dip | Midgefly |  | 4 | 6 |  |  |  |  |  | 10 | 6.55 |
|  | Coleoptera | Coleop | Riffle Beetle | 2 | 2 |  | 8 |  | 2 | 6 | 4 | 24 | 0.52 |
|  |  |  | Water Penny |  |  | 35 |  | 5 | 12 |  |  | 52 | 1.13 |
|  | Odonata | Odon | Damesfly | 79 | 46 | 2 | 56 | 32 | 33 | 37 | 17 | 302 | 6.55 |
|  |  |  | Dragon fly | 5 | 6 |  | 3 | 7 | 10 | 3 | 9 | 43 | 0.93 |
|  | Hemiptera | Hemi | Water strider | 1 |  |  |  |  | 8 |  |  | 9 | 0.20 |
|  |  |  | True Bugs | 1 |  |  | 5 | 2 |  |  | 1 | 9 | 0.20 |
|  |  |  | Water Scorpion |  |  | 15 |  |  |  |  |  | 15 | 0.33 |
|  |  |  | Water Boatman | 25 | 20 |  | 59 |  |  | 25 | 29 | 158 | 3.42 |
| Non-Insecta macroinvertebrates | Gastropoda | Gastr | Gastropods | 539 | 307 | 199 | 332 | 277 | 398 | 190 | 102 | 2344 | 50.8 |
|  | Bivalva (Class) |  | Bivalves | 4 |  |  |  | 2 |  | 1 |  | 7 | 0.15 |
|  | Decapoda | Deca | Shrimps | 14 |  | 5 | 39 | 22 |  | 10 | 45 | 135 | 2.93 |
|  |  |  | Cray fish | 1 | 4 | 2 | 10 | 6 | 2 | 2 | 10 | 37 | 0.80 |
|  | Amphipoda |  | Scuda | 5 |  |  |  |  |  |  |  | 5 | 0.11 |
|  | <u>Euphausiacea</u> |  | Krill | 10 | 4 | 12 | 13 | 4 | 9 | 12 | 14 | 78 | 1.69 |
|  | Oligochaeta | Oligo | Aquatic worms | 79 | 95 | 90 | 190 | 240 | 200 | 135 | 105 | 1134 | 24.6 |
|  | Hirudinea |  | Leeches |  | 2 | 2 | 2 |  |  |  |  | 6 | 0.13 |
| Total |  |  |  | 789 | 521 | 387 | 776 | 622 | 682 | 450 | 378 | 4614 |  |

**Figure 2.**
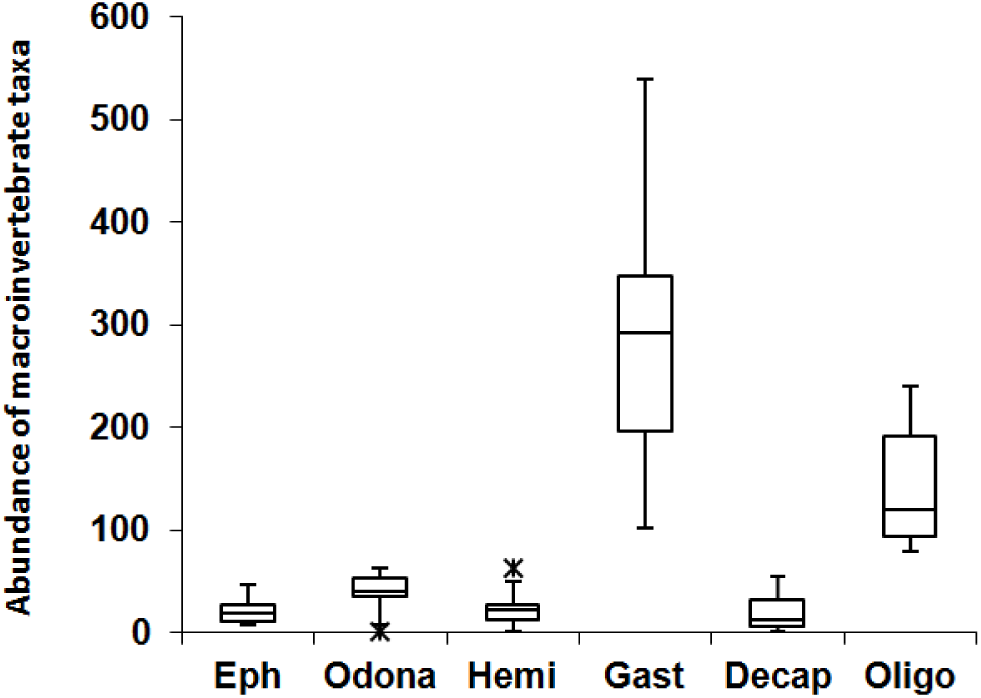
Box plot illustrated the abundance of macroinvertebrate orders, the horizontal line inside each box represents the median, while the top and the bottom of the boxes represent the 25^th^ and 75^th^ percentile, respectively. Vertical lines from the end of the box encompass the extreme point, represent the maximum outlier.

The results of correlation between physicochemical parameters and macroinvertebrates taxa depending on correspondence analysis (CA) were shown in **Fig. (3)**, the first two principal components (D1 = 45.1%, D2 = 21.1%), cumulatively explained 66.2% of the total variance. Generally, Gastropoda and Odonata were clustered together at S1 which distinguished by high chlorophyll-*a* (chl-a) level that indicated to high algal biomass. Most of the Gastropods feed on algae and their increase may be related to an increase in the periphyton and phytoplankton. Algal and diatoms remains to dominate in the gut of snails (Thorp and Covich, 2009). Therefore, gastropods have a greater affinity for nutrient-rich conditions than other macroinvertebrate orders. Additionally, Rosset et al. (2014) stated that local and regional dragonfly (odonata) species richness diversity was not negatively affected by eutrophication except at the local scale in the enriched water bodies.

**Figure 3.**
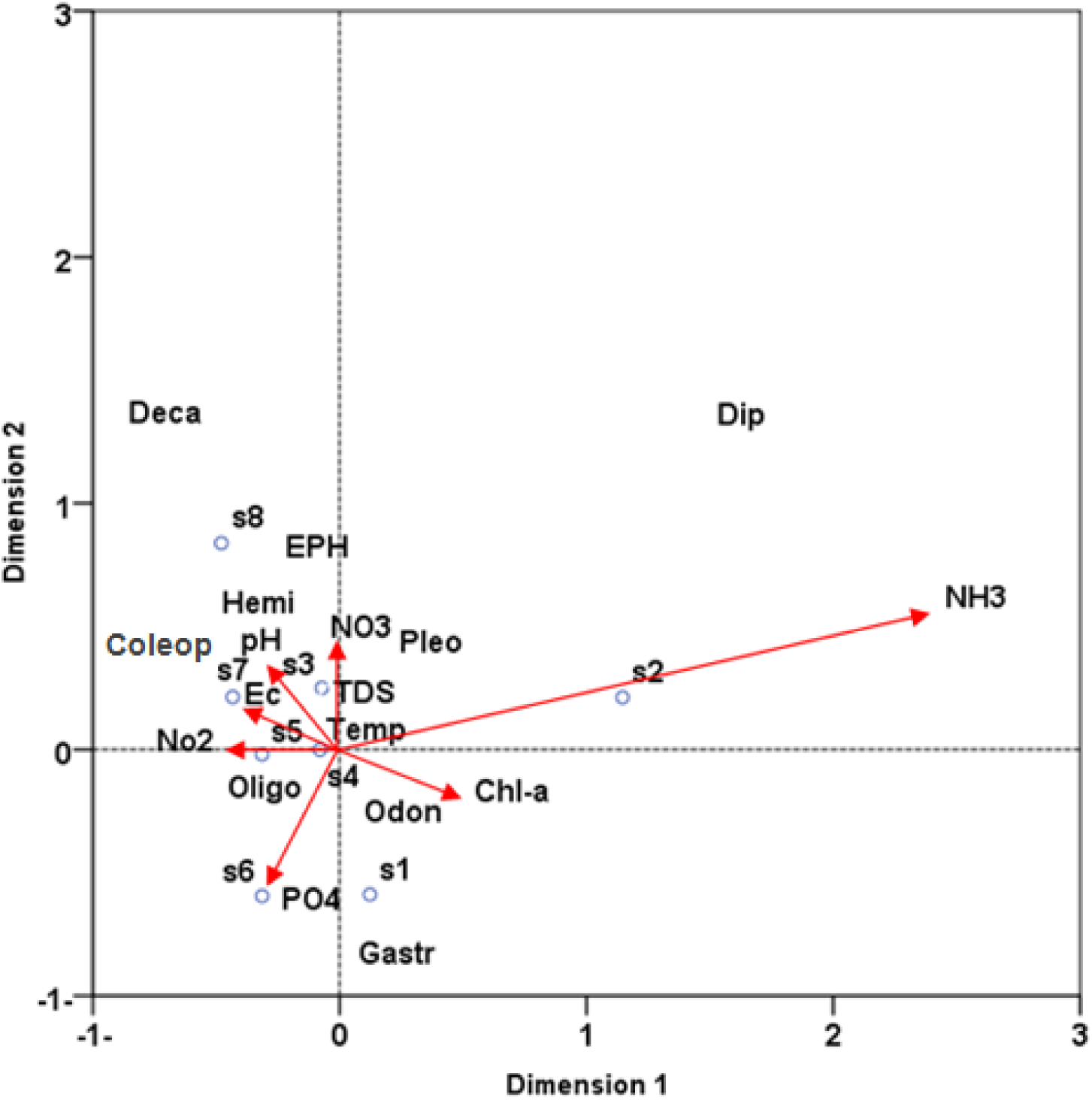
Correlation biplot diagram based on correspondence analysis (CA) of physicochemical variables and macroinvertebrates taxa from investigated stations. Codes of macroinvertebrates were stated in Table 4.

Meanwhile, Oligochaeta was found at S5 and S6 which characterized by high nitrite (NO_2_) and phosphate (PO_4_), respectively. These species are reportedly more abundant in polluted streams and possess the ability to thrive in areas of reduced competition and low concentrations of oxygen (Arimoro et al., 2007). Meanwhile, the rest taxa avoid founding at S2 due to high ammonia (NH_3_) and nitrate (NO_3_).

Regard to the calculated Diversity Index (H′) ranged (1.14 – 2) which indicated that the structure of macroinvertebrates habitat was poor. Also, the calculated Evenness Index (J) ranged (0.016 – 0.043) which indicated that individuals were not distributed equally. In addition, *Hilsenhoff* Biotic Index (HBI) results showed that all examined stations have a significant organic pollution, except station 8 was fairly significant pollution **(Table 5)**. This organic pollution may be attributed to human activities such as farming and municipal wastewater nearby the tested stations. These results in agree with Dahl et al. (2004) and Ojija (2015) who concluded that agricultural activities, washing and bathing alter physico-chemical parameters of the stream and hence changing the abundance of macroinvertebrates as well as the quality of water. In conclusion, the ecological status of River Nile’s water quality around Gizert El-Warrak is poor based on non-taxonomic measurements of algae (chlorophyll-*a*) and biotic indices of macroinvertebrates which reflected the actual conditions of the water quality. Therefore, such measurements may be considered as a vital method that reflects disturbances in aquatic systems for both short-term and /or long-term.

**Table 5.** Macroinvertebrates-based indices Shannon-Wiener diversity index (H′), Pielou evenness index (J) and *Hilsenhoff biotic index* (*HBI*) in the River Nile at Gizert El-Warrak stations.

| Stations | $H'$ index | $J$ index | HBI | Water quality | Degree of pollution |
| --- | --- | --- | --- | --- | --- |
| 1 | 1.24±0.08 | 0.018±0.01 | 6.62 | Fairly poor | Sig. organic pollution |
| 2 | 1.37±0.09 | 0.022±0.02 | 6.72 | Fairly poor | Sig. organic pollution |
| 3 | 1.51±0.10 | 0.026±0.03 | 6.68 | Fairly poor | Sig. organic pollution |
| 4 | 2.00±0.12 | 0.027±0.03 | 6.64 | Fairly poor | Sig. organic pollution |
| 5 | 1.70±0.12 | 0.043±0.11 | 6.83 | Fairly poor | Sig. organic pollution |
| 6 | 1.14±0.10 | 0.016±0.02 | 6.64 | Fairly poor | Sig. organic pollution |
| 7 | 1.59±0.11 | 0.023±0.03 | 6.39 | Fairly poor | Sig. organic pollution |
| 8 | 1.96±0.11 | 0.028±0.02 | 5.99 | Fair | Fairly sig. organic pollution |

## Acknowledgments

The authors express their deep thanks to Dr. Sayeda Mohamed Abdo, Research Water Pollution Department, National Research Centre, Dokki, Egypt, for her help, identified phytoplankton and water chemicals analysis.

